# Topographic diversity of structural connectivity in schizophrenia

**DOI:** 10.1101/282145

**Authors:** Hongtao Ruan, Qiang Luo, Lena Palaniyappan, Wenlian Lu, Chu-Chung Huang, Chun-Yi Zac Lo, Albert C Yang, Mu-En Liu, Shih-Jen Tsai, Ching-Po Lin, Jianfeng Feng

## Abstract

**Background:** The neurobiological heterogeneity of schizophrenia is widely accepted, but it is unclear how mechanistic differences converge to produce the observed phenotype.

**Aims:** Establishing a pathophysiological model that accounts for both heterogeneity and phenotypic similarity is essential to inform stratified treatment approaches.

**Method:** In this cross-sectional diffusion tensor imaging (DTI) study, we recruited 77 healthy controls (HC), and 70 patients with DSM-IV diagnosis of schizophrenia (SCZ). Heterogeneity was assessed by inter-subject similarity and clustering for subgroups. Common feature was established at a system-level as the diversity (or statistically, the variance) of topographic distribution of structural connectivity (SC). Discriminative powers were demonstrated by classifier using topographic diversity as an input feature.

**Results:** We first confirmed the heterogeneity in SC by showing a reduced between-individual similarity in SCZ compared to HC. Moreover, we found it was not possible to cluster patients into subgroups with shared patterns of dysconnectivity, indicating a high degree of mechanistic divergence in schizophrenia. Topographic diversity was significantly reduced in SCZ (*P* = 7.21×10^−7^, *T*_*142*_ = 5.19 [95% CI: 3.37−7.52], Cohen’s *d*=0.91), and this affected 65 of the 90 brain regions examined (False Discovery Rate < 5%). When topographic diversity was used as a discriminant feature for classifying patients from controls, we achieved a classification accuracy of 80.97% (sensitivity 77.14%, specificity 84.42%).

**Conclusions:** This finding suggests highly individualized pattern of structural dysconnectivity underlies the heterogeneity of schizophrenia, but these disruptions likely converge on an emergent common pathway to generate the clinical phenotype of the disorder.

**Declaration of interest:** LP received speaker fees from Otsuka in 2017; LP received unrestricted educational grants from Otsuka Canada and Janssen Canada in 2017. All other authors declare no conflict of interest.

## INTRODUCTION

Schizophrenia is characterised by dysconnectivity in the brain (1), and structural dysconnectivity has been reported by diffusion imaging studies (2–6). Nevertheless, the spatial localisations of white matter (WM) abnormalities have been diffuse (4, 5, 7, 8) (including corpus callosum, cingulate bundle, and uncinate fasciculus, fibres from superior frontal gyrus, superior temporal gyrus, thalamus, the internal capsule, cerebellum, hippocampus, the parietal lobe, and the occipital lobe), with only moderate effect size given by a large sample size (Cohen’s *d* < 0.42) (7). This lack of consistency has often been ascribed to neurobiological heterogeneity of schizophrenia where region-specific pathology results in distinct subgroups of patients (4, 9). While the between-individual variations in the severity of symptom dimensions and cognitive ability appear to be linked to differential WM integrity (10–13), it is important to note that most studies to date have linked distributed WM deficits to the broad clinical phenotype of schizophrenia, rather than hitherto unknown neurobiological subgroups. If aberrant brain connectivity is indeed a neurobiological substrate of the schizophrenia phenotype, then it is likely that these spatially distinct WM changes converge to deviate certain emergent features of WM architecture. In other words, we hypothesized to see a large effect-size deficit in WM architecture at a system-level despite a low degree of between-individual similarity in spatial distribution of WM abnormalities in schizophrenia. We are going to test this hypothesis in this paper.

## METHODS

### Subjects and Image Acquisition

In the current sample (Table 1, Table S1), we had 70 patients with schizophrenia (35 Male, 35 Female, age 42.1 ± 9.4) and 77 age matched healthy controls (33 Male, 44 Female, age 42.1 ± 9.1). The authors assert that all procedures contributing to this work comply with the ethical standards of the relevant national and institutional committees on human experimentation and with the Helsinki Declaration of 1975, as revised in 2008. All procedures involving human subjects/patients were approved by the Institutional Review Board of Taipei Veterans General Hospital. All participants were recruited from the Taipei Veterans General Hospital, and this study was approved by the Institutional Review Board of Taipei Veterans General Hospital. Written informed consent was obtained from all participants. The age of all participants varied from 20 to 55, and the education years varied from 9 to 17. They are all right-handed. The total score of the Positive and Negative Syndrome Scale (PANSS) was over 35 in all patients at the time of scanning. The PANSS scores were low because of the long-term treatment (Table S1). All images were acquired using a 3T MR system (Siemens Magnetom Tim Trio, Erlangen, Germany) at National Yang-Ming University, equipped with a high-resolution 12-channel head array coil. To minimize the head motion during the scan, each subject’s head was immobilized with cushions inside the coil during the scan. A high-resolution anatomical T1-weighted image was acquired with sagittal 3D magnetization-prepared rapid gradient echo (MPRAGE) sequence: repetition time (TR) = 3500 ms, echo time (TE) = 3.5 ms, inversion time = 1100 ms, flip angle = 7°, field of view (FOV) = 256 mm × 256 mm, 192 slices, slice thickness = 1 mm, voxel size =1.0 mm × 1.0 mm × 1.0 mm. The diffusion images gradient encoding schemes include 30 non-collinear directions (according to the minimal energy arrangement of electron distribution) (http://www2.research.att.com/~njas/electrons/dim3) with b-value 1000 s/mm^2^ and 3 non-diffusion weighted image volume. With the consideration of total brain coverage, each volume consisted of 70 contiguous axial slices (thickness: 2 mm) acquired using a single shot spin-echo planar imaging (EPI) sequence (TR: 11000 ms, TE: 104 ms, NEX: 6, Matrix size: 128 × 128, voxel size: 2 mm × 2 mm × 2 mm, matrix size: 128 × 128).

**Table 1.** Demographics and behavioral statistics for schizophrenia and control groups

|  |  | Schizophrenia | Control | Test Statistics |
| --- | --- | --- | --- | --- |
| **Sex (F—female; M—male)** | | 35 F, 35 M | 44 F, 33 M | $\chi^2_1 = 0.75, P = 0.39$ |
| **Age (year)** | | $42.1 \pm 9.4$ | $42.1 \pm 9.1$ | $T_{145} = 0.03, P = 0.98$ |
| **Education (year)** | | $12.8 \pm 2.2$ | $14.2 \pm 2.1$ | $T_{145} = -3.96, P < 0.01$ |
| **Duration (year)** | | $15.8 \pm 9.2$ | / | / |
| **PANSS** | **Positive scale** | $10.1 \pm 3.0$ | / | / |
| | **Negative scale** | $9.8 \pm 2.3$ | / | / |
| | **General Psycho-pathology scale** | $22.0 \pm 3.4$ | / | / |

### Image Preprocessing and Construction of Connectivity Matrix

Brain was extracted from all images using FSL’s brain extraction tool (14, 15). The diffusion-weighted volumes were corrected for eddy-current induced distortions and movements by affine registration to the no diffusion-weighted (B0) image using the diffusion-specific FSL’s EDDY tool (16).

Fiber-tracking was performed by the MRtrix0.2.12 package (http://www.nitrc.org/projects/mrtrix/, (17)). We first extracted the b=0 image and constructed the brain mask. By the MRtrix, we acquired the diffusion tensor parameters and fractional anisotropy (FA) map. Next, the response function of each voxel above FA threshold of 0.7 was estimated from the DWI, and this was used as the input of the constrained spherical deconvolution computation (18). Finally, we performed deterministic fiber-tracking within whole brain using the orientations provided by constrained spherical deconvolution, with an FA cutoff of 0.15, step size of 0.2 mm, minimum length of 10 mm, maximum length of 200 mm, until 2,000,000 fibers were reconstructed for each individual. The AAL template (19) (without parcellations in cerebellum) was registered from MNI standard space into the space of. each subject’s diffusion data by the program Statistical Parametric Mapping (SPM8, http://www.fil.ion.ucl.ac.uk/spm/, (20)). For each subject, each fiber was labelled according to its termination to the AAL template in native space. From these data, we constructed a 90 × 90 structural connectivity (SC) matrix *A*, whose element *A_ij_* was computed as the mean of the FA values of all included fibers that formed the connection between regions *i* and *j* and incorporated the information on the integrity of the interregional white matter connection. The diagonal elements *A_ii_* of *A* was 0 because we only considered the inter-regional connections here.

### Between-individual Similarity of Whole Brain Connectivity

Increased heterogeneity in patients would result in a decrease in between-individual similarity. We estimated the between-individual similarity by calculating the correlations of the whole brain connectivity (i.e., the FA-weighted SC matrix *A*) between subjects of the same group. This resulted in a similarity matrix for each group. This similarity distribution (elements of upper triangular of similarity matrix) was then compared between healthy subjects and patients with schizophrenia by t-test to examine whether the similarity was significantly different in group level.

### Clustering Patients with Schizophrenia

Decreased similarity might be explained by forming of subgroups with distinct patterns of SC in patients with schizophrenia. We tried to identify these subgroups, if any, by applying different clustering methods to the connectivity pattern, *i.e.*, the FA weighted SC between brain regions. Because of the high dimension of 4005 unique structural connections, we utilized two different dimensionality reduction methods to reduce the dimension of features for clustering.

#### Principal components analysis (PCA)

We first selected 740 connections which were significantly different between patients and healthy controls (*P* < 0.05, uncorrected). Then PCA was applied to these connections, and different numbers of components (ranging from 15 to 60 components which explained at least 30% of all variability) were selected as features for clustering.

#### Canonical correlation analysis (CCA)

Following the approach of detecting subgroups in patients (21), the SC was first selected according to their (Spearman’s rank) correlations with the PANSS scales (Positive scale, or Negative scale, or General Psychopathology scale with *P* < 0.005, uncorrected) in patients. Secondly, by CCA, we identified different numbers of canonical components (i.e. linear combinations of those symptom-associated SC) as features for clustering.

To avoid the possibility that the clustering result was affected by the inherent limitations of a particular clustering method, we systematically tested four different combinations between two clustering methods and two features selection approaches, including PCA-based k-means, CCA-based k-means, PCA-based hierarchical clustering and CCA-based hierarchical clustering.

#### K-means clustering

We applied k-means clustering to patients with PCA-selected features (PCA-based k-means) and CCA-selected features (CCA-based k-means). The number of clusters (*k* value) in our analysis ranged from 1 to 6. The optimal number of clusters was determined by the GAP statistics (22), measuring the quality of the resulting clusters.

#### Hierarchical clustering

We also applied a hierarchical clustering method to identify subgroups of patients with PCA-selected features (PCA-based hierarchical clustering) or CCA-selected features (CCA-based hierarchical clustering, which had been successfully applied recently to define neurophysiological subtypes of depression (23)). The results were tested with different cutoffs (a critical inconsistent value for constructing agglomerative clusters) from 0.6 to 1.6 with step 0.01.

### The Stability of K-means Clustering

To assess the stability of k-means clustering, we applied bootstrap to the adjusted Rand index (ARI) (24–26). Firstly, the original patients’ data with PCA-selected features or CCA-selected features were analyzed with k-means for a given cluster number *k* which could be ranged from 2 to 6, resulting in a clustering solution *P_k_* and the *k* cluster centroid locations *C_k_*. Secondly, from all patients, 70 objects were drawn with replacement to yield a bootstrap sample. The bootstrap procedure was repeated 1000 times. Thirdly, every bootstrap sample was analyzed with k-means to get the corresponding *k* cluster centroid locations 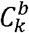 for *b* = 1, 2, …, 1000. Fourthly, 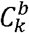 was used to construct the cluster solution 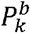 for all patients in the original data by assigning each subject to the cluster with the closest centroid (26). Finally, for each bootstrap, we computed an ARI value with 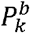 and *P_k_*, and took the meanof 1000 bootstrap ARI values as the final stability *R_k_* for *k*. The scale of *R_k_* helped us to understand the stability of our clustering results and the maximum value of *R_k_* gave us an idea about the number of clusters in patients (25).

### The Topographic Diversity of SC

We computed the variance of the brain SC for each subject as follows:

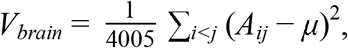

where *μ* was the mean of 4005 structural connections. A larger *V_brain_* indicated a higher global topographic diversity of SC. Similarly, for each brain region defined by the AAL template, the variance of its regional connectivity pattern was calculated to represent the regional topographic diversity for every subject as follows:

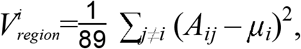

where *μ*_*i*_ was the FA-weighted mean of 89 structural connections connected to region *i*. For group comparison, we implemented a linear regression to compare these variances between patients and controls with age, gender, education used as covariates. To confirm the findings of group differences, we applied 1000 bootstraps to establish 95% confidence intervals of the corresponding statistics. To examine the heterogeneity of regional diversity in patients, we estimated the between-individual similarity by calculating the correlations of the 90 regional diversities between subjects of the same group. This similarity distribution was then compared between two groups by t-test.

### Comparison of Top 10 or Last 10 Connections

For a fixed region *i*, we ranked its FA weighted connections to the other 89 brain region in a descending order. To explore the origins of the group difference in the regional diversity, we calculated the mean FA of the top 10 connections and that of the bottom 10 connections. For each region, the mean value of top 10 (bottom 10) was linearly regressed on age, gender, education and diagnostic group identity (schizophrenia: 0; healthy control: 1) for all participants. After FDR correction for multiple comparisons, this allowed us to test whether the disrupted topographic diversity emerged from aberrations in strong (top 10), rather than weakly connected (bottom 10) WM pathways. The effects of different number of top (bottom) connections range from 5 to 15 were also tested.

### Classification of Patients and Controls

To study if the topographic diversity provided any incremental value in distinguishing patients from healthy controls, we used a random forest classification approach (27). This machine learning approach is particularly useful in this case as it has a built-in cross-validation process to provide ranking of the discriminative features. After the feature selection, another cross-validation procedure was applied to fit the model and evaluate the model performance. Such separation of feature selection and model fitting has been proved to be a promising strategy to prevent over-fitting (28). Two models were built on the basis of: (Model 1. Connectivity model) 4005 SC among 90 brain regions; (Model 2. Combined model) variance of each region, mean FA of the top 10 connections in the regional connectivity pattern, combined with SC.

Firstly, from 4005 structural connections we selected 50 connections as predictors which were most significantly different between patients and controls. For Model 2, we also chose 50 brain regions whose regional diversity differed most obviously between two groups, and used the regional diversity and the mean FA of top 10 connections of those regions as the predictors. Secondly, we ran the random forest algorithm for each model with 2000 trees for 200 times on the predictors to obtain a median importance score for each predictor. The highest 6-15 predictors were used in the final model to classify the subjects into patients and controls. Notably, this technique utilizes bootstrapped cross-validation to reduce overfitting when generating permutation importance scores. Thirdly, using five-fold cross-validation, we compared the classification accuracies of these two models by McNemar’s test (29), and also estimated an accuracy of classification for each model. The results were validated by exploring different numbers of selected predictors for these models.

## RESULTS

### Heterogeneity Observed in SC Patterns of Patients

Compared with healthy controls, patients had decreased between-individual similarity of SC (*P* = 2.82 × 10^−64^, *T_145_* = 17.16, Cohen’s *d* = 0.47), indicating notable within-group heterogeneity among patients (Figure 1A).

**Figure 1.**
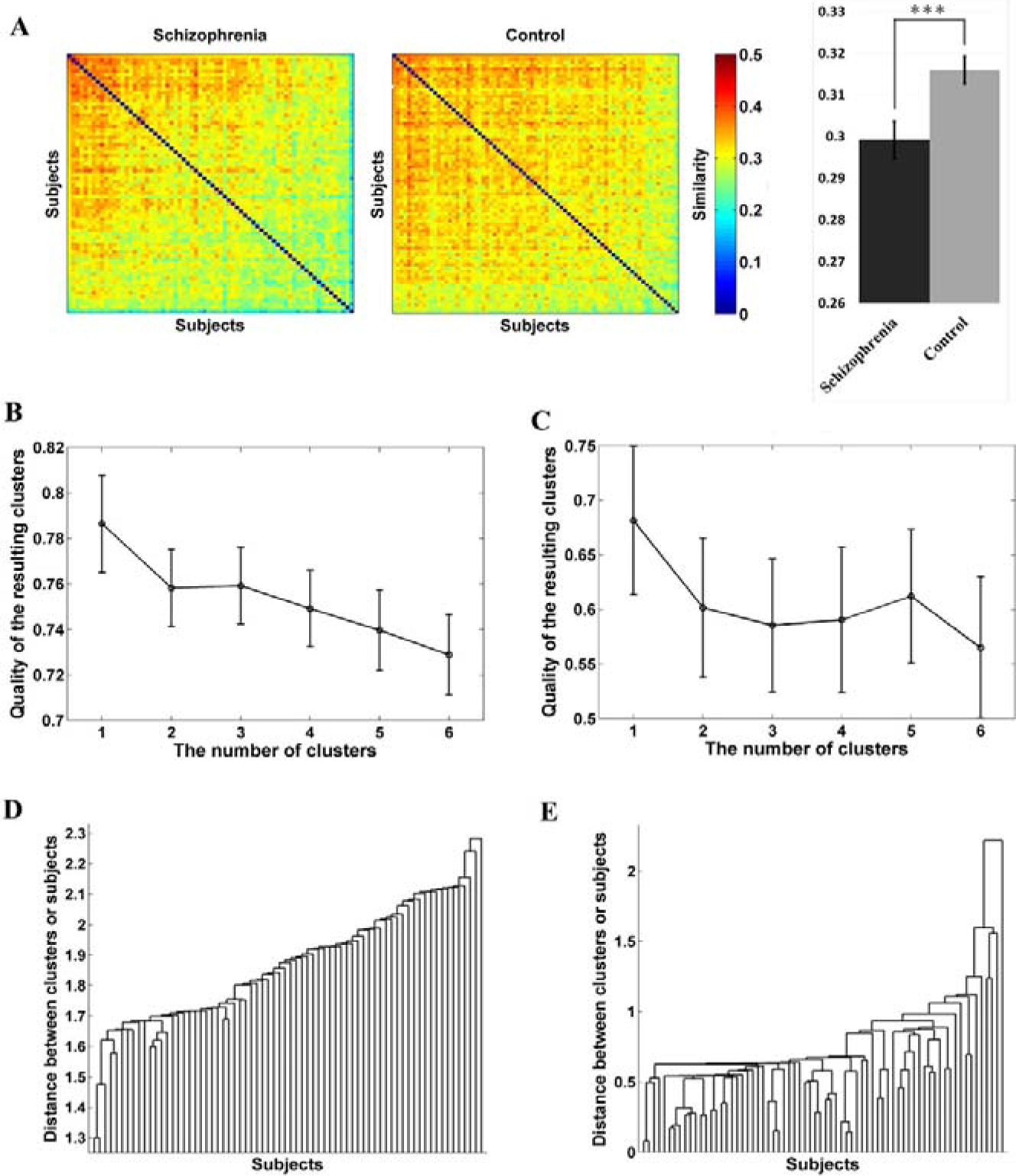
Higher heterogeneity of SC in patients resulted in no robust subgroups. A) Between-individual similarity matrices of two groups, and group comparison of the between-individual similarity; B) The quality of the resulting clusters measured by the GAP statistics for PCA-based k-means clustering, with 50 principal components as features and k = 1, 2, 3, 4, 5 and 6; C) The quality of the resulting clusters measured by the GAP statistics for CCA-based k-means clustering, with three canonical components and k = 1, 2, 3, 4, 5 and 6; C) PCA-based hierarchical clustering for patients using 50 principal components. D) CCA-based hierarchical clustering for patients using three symptom-associated canonical components of the SC.

Furthermore, we found SC patterns in patients were too heterogeneous to detect any distinct SC pattern shared by any subgroup of patients, as no subgroup could be clearly identified by neither k-means nor hierarchical clustering. According to the quality of clusters given by the GAP statistic for k-means (22), we found that the optimal number of clusters was 1 in patients (Figure 1B-C; Figures S1), despite systematically varying of the complexity of feature space (principal components or canonical components; Tables S2 and S3). According to the stability of clusters assessed by 1000 bootstraps of adjusted Rand index (ARI), we found all clustering results had low stabilities (<0.77) (Table S4 and S5). Similarly, no cluster could be clearly defined by hierarchical clustering as shown more intuitively in dendrograms (Figure 1D-1E; Figures S2).

### Topographic Diversity of SC Decreased More Homogenously in Patients

Compared with patients, HC had greater diversity in the whole brain SC (*P* = 7.21 × 10^−7^; *T_142_* = 5.19 [95% CI: 3.37-7.52]; Cohen’s *d* = 0.91), indicating that the difference between strong connections and weak connections were larger in HC (Figure 2A). In contrary to SC, patients compared with HC became more similar to each other in terms of topographic diversity as they had increased between-individual similarity of topographic diversity (*P* = 4.66 × 10^−11^, *T_145_* = −6.60, Cohen’s *d* = 0.18) (Figure 2B).

**Figure 2.**
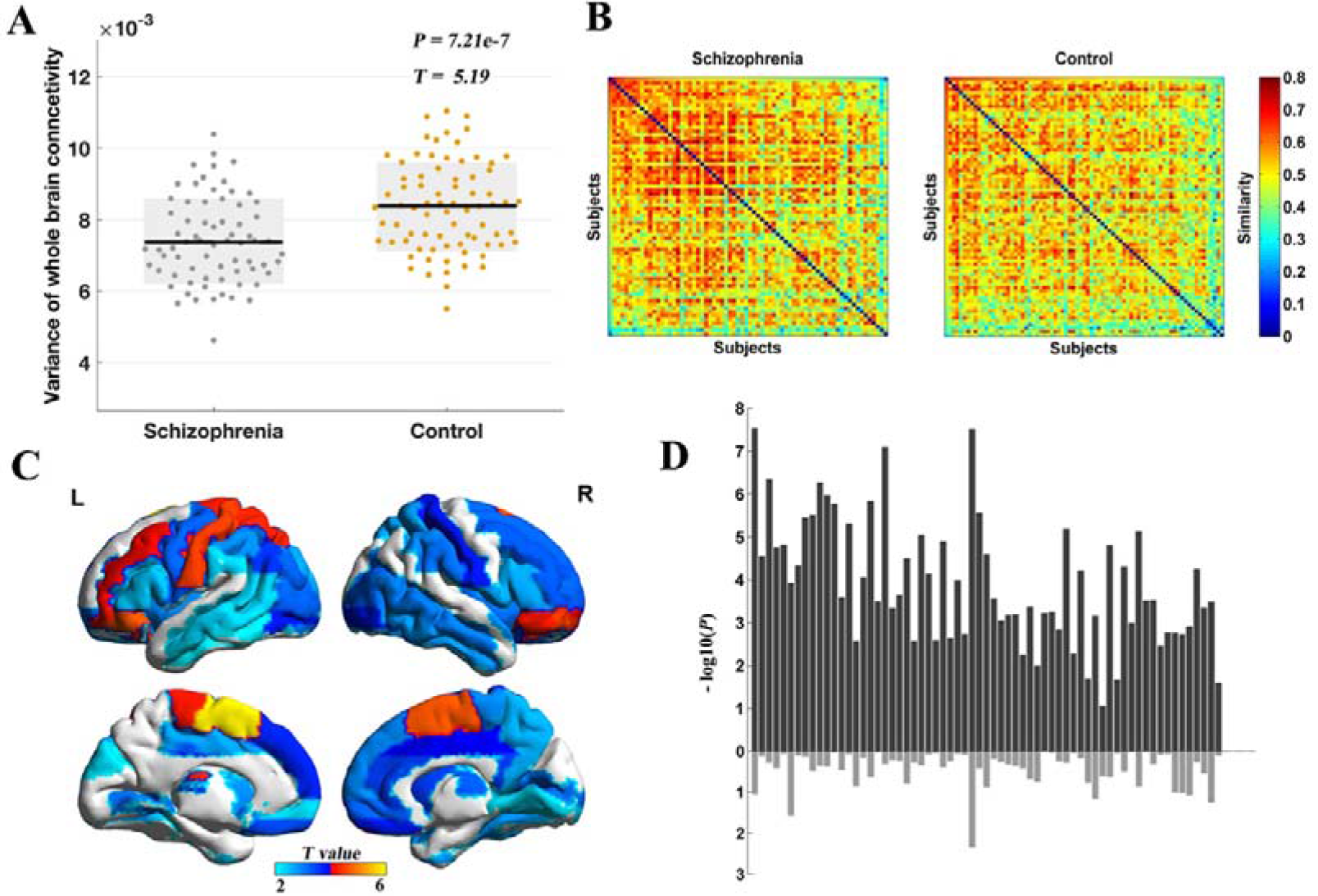
Topographic diversity of brain structural connectivity reduced in patients. A) Comparison of the within-individual variance in the whole brain SC (structural connectivity); B) Between-individual similarity matrices of regional diversity in two groups; C) Spatial distribution of the 65 brain regions with significantly reduced variances in their regional connectivity patterns in patients compared with controls; D) The *P* value of between-group comparison for the mean FA of both the top 10 (upper part in the figure) and bottom 10 (lower part in the figure) connections in the regional connectivity pattern of these 65 brain regions, the abscissa axis represented brain regions and the vertical axis was −log_10_(*P*).

After FDR correction, we found that the regional diversity of 65 brain regions (72.2% of 90 brain regions) were reduced in patients compared with controls (Figure 2C). Areas showing reduced diversity were mainly located in frontal lobe, prefrontal lobe, parietal lobe, occipital lobe and subcortical areas. The top 5 most abnormal regions were SMA.L (*P* = 2.55 × 10^−8^; *T_142_* = 5.90 [95% CI: 3.90-8.00]; Cohen’s *d* = 1.04), SMA.R (*P* = 6.04 × 10^−6^; *T_142_* = 4.70 [95% CI: 2.83-6.89]; Cohen’s *d* = 0.83), ORBmid.R (*P* = 1.22 × 10^−5^; *T_142_* = 4.53 [95% CI: 2.43-6.42]; Cohen’s *d* = 0.80), ORBinf.L (*P* = 5.08 × 10^−6^; *T_142_* = 4.74 [95% CI: 2.73-6.92]; Cohen’s *d* = 0.84) and PoCG.L (*P* = 1.26 × 10^−5^; *T_142_* = 4.53 [95% CI: 2.59-6.26]; Cohen’s *d* = 0.80).

Furthermore, we found for 64 out of 65 brain regions identified above, the connectivity strength of the top 10 strongest connections but not the bottom 10 weakest connections (*P* < 0.05 FDR corrected) were reduced in patients (Figure 2D; Table S6). We also confirmed this finding using different number of top (bottom) connections ranging from 5 to 15.

Current antipsychotic dose equivalents had no association with the regional or the global diversity (Table S7).

### Topographic Diversity Improved the Classification between Patients and Controls

The accuracy rate of Model 1 was 73.45% (sensitivity 72.86% and specificity 74.03%) with nine connections, predominantly frontal (Table S8), selected by the random forest. Introducing the topographic diversity to the model the accuracy rate increased to 80.97% (sensitivity 77.14% and specificity 84.42%) in Model 2. Such improvement of classification accuracy was significant (*P* = 0.01, χ_1_ = 1.34 by McNemar’s test (29); Figure 3A). The effects of topographic diversity in classification were consistently positive when varying the number of predictors selected (Figure 3B).

**Figure 3.**
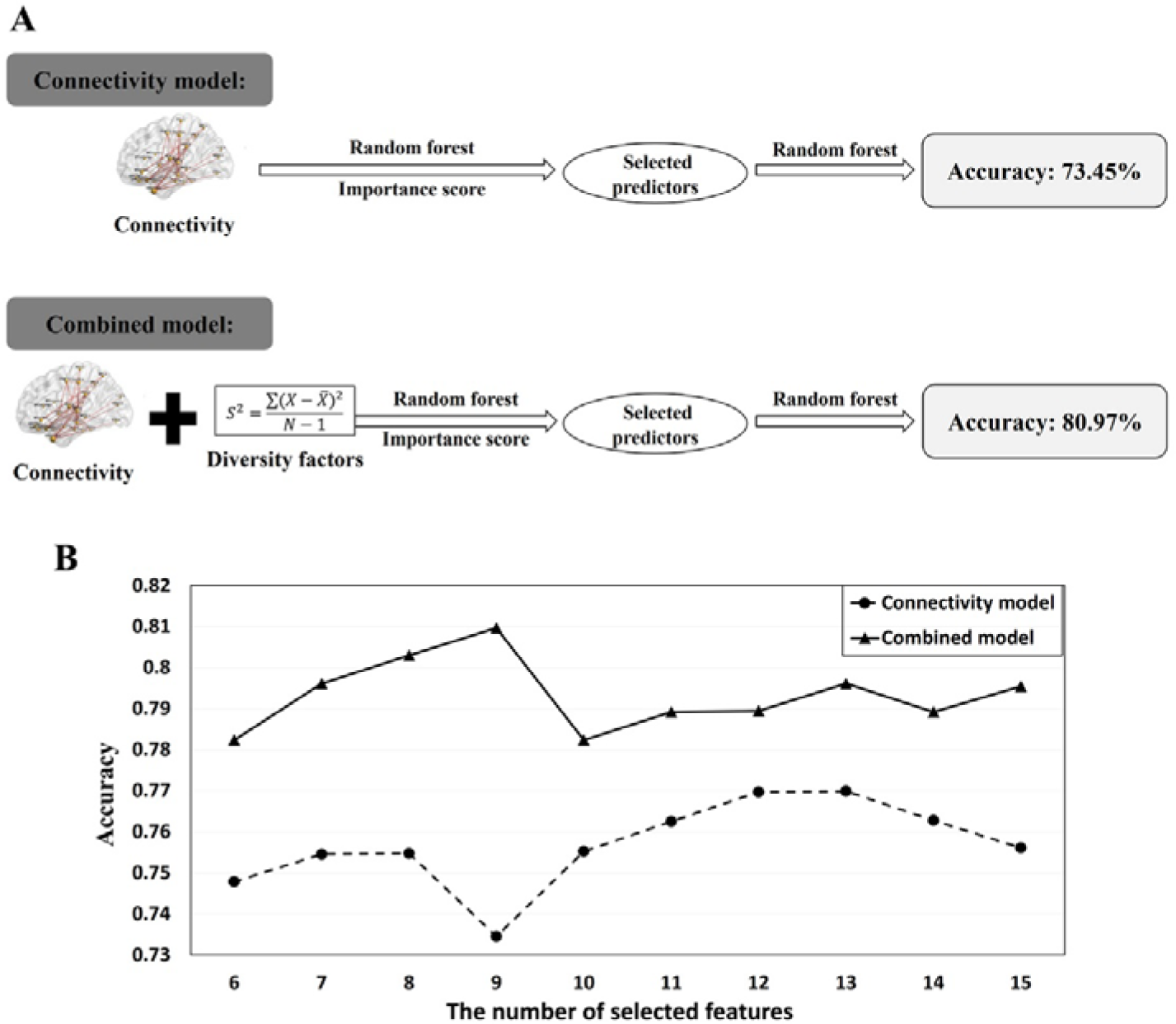
Topographic diversity as a system-level feature improved the classification accuracy between patients and controls. A) Two models with different sets of features and the accuracies; B) The accuracies of two different models with the number of selected features ranging from 6 to 15.

## DISCUSSION

To the best of our knowledge, this is the first study to systematically examine if the individually distinct WM changes seen in patients with schizophrenia lead to a convergent effect that is relevant to the expressed clinical phenotype. We report 3 major findings from the current study. 1) The between-individual spatial similarity in WM connectivity is greatly reduced in schizophrenia compared to healthy controls. 2) Despite the spatial dissimilarity, patients with schizophrenia, as a group, show large effect-sized reductions in topographic diversity, which when used in conjunction with the observed pattern of regional changes in each patient, results in successful discrimination of the patients from controls. 3) The reduced topographic diversity relates almost exclusively to disruptions in strongly connected WM pathways.

Our findings add an important clarification to the divergent results of DTI studies in schizophrenia. In patients with schizophrenia, subtle disruptions in WM connectivity occur in a person-specific (idiosyncratic) fashion. The patterns of resulting deficits, when combined, are sufficiently discriminative of the phenotype, but this is not due to a generalized non-specific deficit affecting all WM pathways. Our results suggest that the WM deficit in schizophrenia is likely to be specific to the strongly connected pathways that are more liable to have direct axonal connections. This inference reconciles the spatial divergence (30) with the discriminatory accuracy*(31–34)* (generally 70-90%) when using multivariate approaches in the DTI studies.

We lacked neuropsychological data to examine cognitive specificity of regional WM aberrations. Interestingly, there was no discernible association between topographic diversity and symptom burden, but we observed a notable canonical relationship between symptom severity and regional WM aberrations. This supports the notion that the highly variable, small magnitude, lower-level mechanistic markers may operate to produce individual symptoms, while the system-level indicators are more relevant to the emergence if the broader clinical phenotype of schizophrenia. In this sense, topographic diversity may represent the elusive neural integrative deficit (35) that is proximal to clinical phenotype but relatively distal to etiological factors (36).

There is an enthusiastic acceptance that schizophrenia is such a disorder with neurobiological heterogeneity; but the evidence demonstrating such heterogeneity is not yet available, or at best, weak (37). In most studies, the absence of clear demarcation of a neurobiological feature between patients and controls (*i.e.*, small effect size) or the demonstration of increased variance (38) among patients is taken to be supportive of the underlying heterogeneity. While this is certainly a viable explanation, we report that convergent neurobiological abnormalities at a systemic level can stem from diverse changes. Thus, the neurobiological heterogeneity inferred at a lower level, may not be observable at the ‘systems’ level. It is important to note that the lack of clustering solutions does not negate the neurobiological heterogeneity of schizophrenia. On the contrary, these results are expected if there is a very high level of heterogeneity. But the large effect-size reduction in systems-level property (diversity), reminds us that it may be premature to dismiss the possibility that schizophrenia, as observed by contemporary clinicians, could be a system-level deficit in WM architecture that may vary only in degree across individuals (Figure S3 summarizes the possible mechanistic nature of schizophrenia).

### Limitation

It is important to consider 2 important caveats in this study. We recruited a sample of medicated patients with an established history of schizophrenia. While this reduced the heterogeneity that is typically seen in early stage, drug-naïve samples with relatively weaker diagnostic stability, a selection-bias towards a single group of patients with relatively more severe illness and poor prognosis cannot be excluded. It is rather surprising that this bias has not improved the between-individual similarity in WM aberrations. Secondly, compared to recent consortium-based neuroimaging studies (7), our sample size was relatively modest. This might explain the lack of clustering solutions within the patient group. Nevertheless, it is acknowledged that the heterogeneity in psychiatric disorders are expected to be diffuse (39); as a result, an ‘optimal’ clustering solution may not be achieved even in large samples (40).

### Conclusion

In summary, the convergence of highly individualized (idiosyncratic) pattern of structural dysconnectivity resulting in reduced topographic diversity in schizophrenia validates the century-old concept of dysconnectivity or integration deficit as a binding feature that brings together seemingly disparate dimensions of symptoms in patients with schizophrenia. When parsing the heterogeneity of psychosis, it is critical to accommodate emergent system-level properties that can capture the gestalt of the currently used clinical construct of schizophrenia.

## Acknowledgement

This work was supported by grants (V105C-008, V105E17-002-MY2-1) from the Taipei Veterans General Hospital. QL is supported by the grants from the National Natural Science Foundation of China (No. 11471081 and 81873909) and Natural Science Foundation of Shanghai (No. 17ZR1444400). C.P.L was supported in part by funding from Ministry of Science and Technology (MOST 104-2633-B-400-001, MOST 104-2218-E-010-007- MY3, MOST 104-2221-E-010-013), National Health Research Institutes (NHRI-EX-10310EI) and Academia Sinica (AS-104-TP-B10) of Taiwan. A.C.Y was supported by Taiwan (MOST 101-2314-B-075-041-MY3, 104-2314-B-075-078- MY2). S.J.T. was supported by Taiwan (MOST 103-2314-B-075-067-MY3, MOST 104-2745- B-075-002). JF is partially supported by the key project of Shanghai Science & Technology Innovation Plan (No. 15JC1400101 and No. 16JC1420402) and the National Natural Science Foundation of China (No.71661167002 and No. 91630314). The research was also partially supported by the Shanghai AI Platform for Diagnosis and Treatment of Brain Diseases (No. 2016-17). The research was also partially supported by Base for Introducing Talents of Discipline to Universities (No.B18015). JF was a Royal Society Wolfson Research Merit Award holder. LP is supported by the Academic Medical Organization of Southwest Ontario; Bucke Family Fund; Chrysalis Foundation and Canadian Institute of Health Research Foundation Grant.

